# A Reappraisal of the Role of the Mammillothalamic Tract in Memory Deficits Following Stroke in the Thalamus

**DOI:** 10.1101/2025.06.13.659501

**Authors:** Julie P. C. Vidal, Lola Danet, Jérémie Pariente, Jean-François Albucher, Patrice Péran, Emmanuel J. Barbeau

**Affiliations:** Brain and Cognition Research Center (UMR 5549), CNRS, University of Toulouse, Toulouse, France; Toulouse Neuroimaging Center (UMR 825), Inserm, University of Toulouse, Toulouse, France; Neurology Department, Toulouse University Hospital Purpan, Toulouse, France

**Keywords:** Thalamus, mammillothalamic tract, memory, thalamic stroke, connectivity

## Abstract

The thalamus, traditionally viewed as a sensory relay station, is now recognized for its critical role in higher-order cognitive functions, including memory. While most research has focused on its distinct nuclei, the thalamus also contains crucial white matter tracts, such as the mammillothalamic tract (MTT), part of the Papez circuit. Despite its established role in memory, the MTT remains underappreciated in studies on the cognitive role of the thalamus, increasing the risk of misattributing memory functions to nearby thalamic nuclei. This study investigates the memory impact of thalamic strokes by considering disruption to the MTT.

We examined 40 patients with chronic ischemic thalamic lesions and 45 healthy controls using neuropsychological assessments, focusing on the Free and Cued Selective Reminding Test (FCSRT) and high-resolution structural neuroimaging. Advanced imaging techniques, such as symptom mapping and disconnectome analyses, were employed to analyze the relationships between lesion sites, tract disconnections and cognitive outcomes. Additionally, the expression of calbindin-rich matrix cells and parvalbumin-rich core cells was examined to assess how the connectivity property of thalamic cells, and its disruption, can relate to cognitive deficits following a thalamic stroke.

Patients with left-sided thalamic lesions, especially those involving the MTT, showed significant memory impairments. Symptom mapping identified a specific cluster of lesioned voxels in the left anterior-lateral thalamus, including the MTT, as being strongly associated with poorer memory performance. Disconnectome analysis confirmed that verbal memory deficits were associated with disruption of the MTT. Furthermore, a striking overlap was observed between the critical regions linked to memory deficits and calbindin-rich thalamic areas. However, this calbindin-rich region was also disrupted in patients without memory impairment, revealing that MTT disruption, and not lesions to this region, induced memory deficits in these patients. The identified FCSRT deficit cluster overlapped with brain regions consistently linked to memory processes following fMRI meta-analytic mapping of memory-related keywords.

These findings challenge the traditional focus on thalamic nuclei and connector hubs, indicating a need to reappraise the importance of the MTT in memory impairment after a thalamic stroke. This study advocates for a shift in thalamic research, emphasizing the need to investigate tract-specific contributions and disruptions of the MTT but also of other tracts such as the interthalamic adhesion or amygdalo-fugal pathway, rather than holding only to a “nuclei-centric” view of the thalamus.

## 1. Introduction

The thalamus has long been considered a key relay station for sensory transmission to the cortex. However, its critical role in higher-order cognition, including memory, executive functions and language, is now also widely acknowledged (Sherman, 2016). With its extensive cortical and subcortical connections, the thalamus facilitates brain communication through its diverse nuclei. This has led to a “nuclei-centric” view of the thalamus. Nevertheless, various white-matter tracts, such as the mammillothalamic tract, the interthalamic adhesion and the amygdalofugal pathway, also cross the thalamus and must be considered. This is especially important when the thalamus is damaged, as there is a risk of mistakenly attributing functions to nuclei that actually depend on these tracts.

Of these white matter tracts, the mammillothalamic tract (MTT, Fig. 1A) is pivotal for memory. Early hypotheses, such as those by Aggleton and Brown (1999, 2006), suggested that the MTT mediates episodic recall within the Papez circuit or extended hippocampal circuit (Fig. 1B, 1C). Clinical and tractography studies support this role in episodic memory, linking severe amnesia in thalamic stroke patients to MTT damage (Van der Werf *et al*., 2003; Yoneoka *et al*., 2004; Carlesimo *et al*., 2007; Cipolotti *et al*. 2008; Kwon *et al*., 2010; Danet *et al*., 2015). Recent research further associates the MTT, along with the mammillary bodies and anterior thalamus, with contextual memory. This involves processing the “what” and “where” aspects of episodic experiences. These same structures are also involved in the theta rhythm essential for coordinating brain regions involved in memory (Dillingham *et al*., 2015; 2021; Aggleton *et al*., 2023).

**Figure 1:**
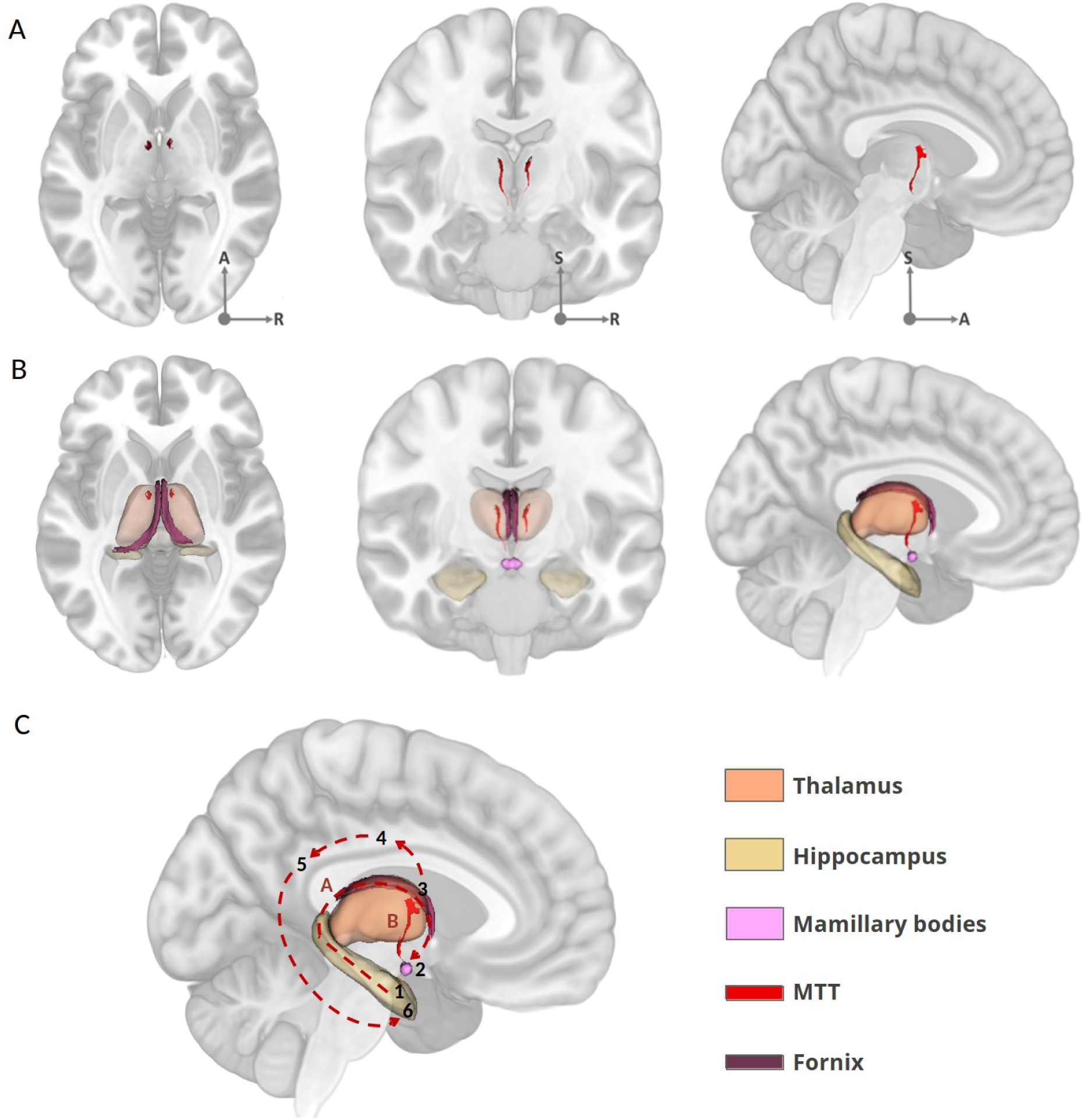
A- The MTT (in red), displayed on axial, coronal and sagittal slices of the MNI152 template. B- The MTT (in red) and the fornix (in purple), displayed on axial, coronal and sagittal slices of the MNI152 template along with other key structures of the Papez circuit. C- 3D representation of the Papez circuit on a sagittal MNI152 slice along with parcellations of subcortical structures and key tracts. The fornix (A, dashed line overlapped with a red manual segmentation) transfers information from the hippocampus (1) to the mammillary bodies (2), which are connected to the anterior thalamic nuclei (3) via the MTT (B, red manual segmentation). The anterior nuclei relay this information to the cingulate gyrus (4), which is connected to the posterior cingulate (5), ultimately transferring the input back to the parahippocampal gyrus (6). Subcortical segmentations were generated either from FreeSurfer or were manually segmented (MTT, Mammillary bodies, Fornix). A: Anterior, S: Superior, R: right.

Variability in thalamic segmentation methods (e.g., HIPS-THOMAS: Su *et al*., 2019; Vidal *et al*., 2024; Morel’s atlas: Morel, 2007; Krauth *et al*., 2010) can sometimes result in inconsistencies in MTT characterization. Because thalamic segmentation can be approximate, depending on resolution, contrast and registration techniques (Segobin *et al*, 2024), direct visual assessment is preferable. The course of the MTT can now be traced relatively easily within the thalamus using high-resolution, high-quality, T1w MRI sequences (Fig. 1C). This makes it simpler to directly assess whether the MTT is damaged or preserved, whereas most previous studies relied on inference from the literature and from atlases. This advance calls for a reappraisal of the MTT’s role in memory through direct assessment of its structural integrity on MRI (Danet *et al*, 2015). In addition, and despite compelling evidence of its role in memory, the MTT often, and surprisingly, remains overlooked in studies on thalamic lesions and related pathologies, probably because of the “nuclei-centric” view of the thalamus. This oversight can lead to biases in attributing memory deficits to specific thalamic nuclei or the thalamus as a whole, rather than to this tract (e.g. Jong *et al*., 2008; Low *et al*., 2019; Scharf *et al*., 2020; Hwang *et al*., 2021).

With advancements in neuroimaging, contemporary research has however shifted from focusing on individual thalamic nuclei to thalamic hubs: regions that integrate widespread cortical and subcortical networks. Hwang *et al*. (2021) identified a critical thalamic region linked to diverse cognitive deficits. This region is rich in calbindin matrix cells, which facilitate inter-network connectivity, an essential feature of connector hubs. Those cells are distinct from parvalbumin-rich core cells, which are involved in intra-network connectivity for specialized cognitive functions, similar to provincial hubs (Jones, 2001, 2009; Gratton *et al*., 2012; Hwang *et al*., 2017). This thalamic hub encompasses the anterior, mediodorsal, ventral medial, intralaminar, ventral anterior and ventral lateral nuclei. Despite methodological differences, this thalamic hub overlaps across multiple studies, suggesting a crucial area for memory and other high-level cognitive domains. However, the MTT also runs through this hub, although this has not been specifically assessed. In light of these recent findings, another reason to reappraise the role of the MTT is to determine whether memory impairment following damage to this hub region is related to the hub itself or to the MTT.

Given the variability in cognitive outcomes following isolated thalamic lesions, there is an increasing need to update our understanding of the relationship between the MTT, the thalamus and memory. This study investigated memory in a large cohort of chronic ischemic thalamic stroke patients. By combining high-resolution imaging with advanced symptom mapping, this research aimed to clarify the role of the MTT in memory.

## 2. Materials & Methods

### 2.1. Participants

This cohort study involved 45 healthy participants (ages 20–69, mean age 48.5, 20 males) and 40 patients (ages 23–75, mean age 51.1, 25 males) with isolated chronic ischemic thalamic lesions (28 left, 6 right, 6 bilateral). Participants were prospectively recruited across two studies from the stroke units of Toulouse hospitals between 2012–2013 and 2019–2021. The first study, approved by the Institutional Review Board “Comité de Protection des Personnes Sud-Ouest et Outre-Mer no. 2-11-0,” (committee for the protection of subjects of biomedical research, south-west and overseas) included 20 healthy individuals and 20 patients under the age of 80 who had experienced a thalamic stroke, with three cases of thalamic lesions marginally extending outside the thalamus (Danet *et al*., 2015). The second study, authorized by the “Comité de Protection des Personnes Ile-de-France IV,” involved 20 healthy individuals and 20 patients under 70 years old, all with at least one stroke lesion in the dorsomedial nucleus and no extra-thalamic damage. Both studies required all participants to have a Fazekas and Schmidt score of ≤ 2. Inclusion criteria for both cohorts included the detection of a first symptomatic thalamic infarct, regardless of prior neurobehavioral complaints, and the absence of known neurovascular, inflammatory or neurodegenerative diseases. All participants underwent clinical, neuropsychological and neuroimaging evaluations on the same day. For patients, these assessments were performed at least three months post-stroke (median: 501 days, range: 91 to 2674 days). Informed consent was obtained from all participants.

### 2.2. MRI Acquisition

For the first 20 patients and 20 healthy subjects, 3D T1-MPRAGE sequences were acquired on a 3T scanner (Philips Achieva) with the following parameters: 1*1*1 mm voxel size, TE = 3.7 ms, TR = 8.2 ms, flip angle = 8°, FOV = 240*240, spacing between slices = 0 mm. 3D T2-FLAIR (Fluid Attenuated Inversion Recovery) sequence parameters were: 1*1*1 mm voxel size, TE = 338 ms, TR = 8000 ms, TI = 2400 ms, FOV = 240*240, spacing between slices = 0 mm. For the next 20 patients and 25 healthy subjects, 3D T1-MPRAGE sequences were acquired on a 3T scanner (Philips Achieva) with the following parameters: 0.9*0.9*1 mm voxel size, TE = 8.1 ms, TR = 3.7 ms, flip angle = 8°, FOV = 256*256, spacing between slices = 0 mm. 3D T2-FLAIR sequence parameters were: 1*1*1 mm voxel size, TE = 343ms, TR = 8000ms, TI = 2400ms, FOV = 240*240, spacing between slices = 0 mm.

### 2.3. Neuropsychological Assessment

The Free and Cued Selective Reminding Test (FCSRT, Van der Linden *et al*., 2004; Grober *et al*., 1988; Grober *et al*., 2000) is a widely used verbal memory test designed to assess anterograde episodic memory, specifically evaluating processes like encoding, retrieval, storage and consolidation of information.

The FCSRT involves learning a list of 16 words, and the following test stages:

- Immediate Recall: Immediate cued recall follows each word presentation on a card, where the participant recalls each word with the help of a category cue.
- Free Recall Sum: After a brief distraction task (e.g., counting backward for 20 seconds), the patient attempts to recall all 16 words in any order for two minutes. This procedure is repeated two more times to reinforce memory consolidation. The total number of words recalled is recorded and summed for the three trials. This measures the participant’s spontaneous recall ability.
- Free Recall Total: If the patient fails to recall a word during a free recall phase, the examiner provides a semantic cue (category) to prompt recall (cued recall). The combined total of words remembered through free and cued recall produces a total recall score. Scores from these repeated trials give an overall assessment of the patient’s ability to retain and retrieve information over multiple attempts and the impact of cues on recall accuracy.
- Delayed Free Recall: After 20 minutes, the patient is asked to perform a free recall of the 16 words.
- Delayed Recall Total: Combining delayed free recall and cued.

In summary, each phase of the FCSRT— encoding with cues — immediate, cued and delayed recall— provides insights into different aspects of memory function. The structured cues and repeated recall trials allow researchers and clinicians to gauge how effectively the individual can encode and retrieve memories, making the FCSRT a comprehensive tool for memory assessment.

To control for language performances, semantic and literal fluency tests (Godefroy, 2008) as well as the ExaDé confrontation naming test (Bachy-Langedock, 1989) were used.

### 2.4. Statistical Analyses

Comparisons were performed using a χ² test for nominal data. Participants’ FCSRT scores were standardized to z-scores based on normative data. A Kruskal-Wallis test was used for quantitative group comparisons, followed by Bonferroni-corrected Dunn’s tests to identify specific subgroup differences. A Wilcoxon test was used to compare two groups, along with an effect size assessed by a Rank-Biserial correlation. Results were presented as boxplots (median ± interquartile range) using z-scores by group to analyze individual performance.

### 2.5. Neuroimaging Analyses

#### 2.5.1. MTT and Papez circuit illustration

To illustrate the Papez circuit and the MTT, the MNI152 template was segmented using FreeSurfer 7.1 subcortical parcellation (Saygin *et al*., 2017). Additionally, manual segmentation of bilateral MTT, fornix and mammillary bodies was generated on MRIcron (J.P.V.) (Rorden & Brett, 2000). Finally, the figure was rendered in 3D using 3D Slicer software, providing a comprehensive circuit visualization. The MTT is visually identifiable in each T1-weighted MRI view (Fig. 1).

#### 2.5.2. Lesion Characterization and Location

Lesions were manually segmented on the native T1-weighted (T1w) images by two independent investigators (J.P.V and L.D) using MRIcron, and the resulting overlapping mask was used for further analysis (mean Dice coefficient = 0.84). This threshold was arbitrarily chosen to account for potential registration inaccuracies during segmentation and to exclude very small lesions unlikely to compromise the tract’s structural integrity. Notably, because the MTT is a thin white matter bundle that is typically only segmented in the portion traversing the thalamus—and may be as narrow as a single voxel on axial slices—even minimal damage can result in apparent tract interruption. Therefore, a threshold of 3 lesioned voxels was set to avoid overestimating disruption due to single-voxel noise or minor lesions, while still capturing cases likely to affect the MTT’s connectivity. Disruption was assessed using Morel’s segmentation method, which enables quantification of lesioned voxels within the thalamic portion of the MTT (Morel, 2007; Krauth et al., 2010). This approach ensures consistent identification of true tract involvement.

#### 2.5.3. Normalization

Native T1w images and segmented lesions were registered to the MNI152 template using affine and diffeomorphic deformations for bilateral lesions and enantiomorphic transformations for unilateral lesions, following skull stripping with the BCBtoolkit (Foulon *et al*., 2018). Normalized segmented lesion masks were used to create an overlap map of patients’ lesions, categorized by infarct laterality, on the MNI152 template using MRIcroGL.

#### 2.5.4. Disconnectome

To map the impact of thalamic strokes on white matter tracts, the Disconnectome Maps software from the BCBtoolkit was used (Forkel *et al*., 2014). For each patient, the software used the normalized segmented lesion as a seed for tractography in 10 healthy subjects, generating an output map displaying voxel-wise probabilities of disconnection within the MNI152 space. The probability of disconnection for each tract was determined using the Tractotron software from the same toolkit. Tracts with a disconnection probability > 50% were considered significantly affected (Foulon *et al*., 2018) and used as the disconnectome map threshold. This map was binarized for symptom mapping analysis.

#### 2.5.5. Symptom mapping

The effect of lesions and structural disconnection on FCSRT subtest z-scores and, as a control, language test performance, were assessed using non-parametric 2-sample t-tests in Randomise (Winkler *et al*., 2014; https://fsl.fmrib.ox.ac.uk/fsl/fslwiki/Randomise/UserGuide). The analysis used a 4D file containing patient lesion segmentations or disconnectome maps as input with 5,000 permutations, family-wise error correction, and lesion volume as a covariate (Souter *et al*., 2022). To enhance statistical power, only lesioned voxels shared by at least three patients were included, reducing multiple comparisons. The output consisted of lesioned or disconnected voxel clusters linked to poorer subtest performance, represented as t-value maps thresholded to retain only significant voxels. These voxel clusters were displayed as a t-map in MNI152 space, overlaid with HIPS-THOMAS segmentation (Histogram-based Polynomial Synthesis-Thalamus Optimized Multi-Atlas Segmentation; Su *et al*., 2019; Vidal *et al*., 2024) to identify the affected thalamic nuclei.

### 2.6. Relative concentration of core cells and matrix cells

Following Hwang *et al*. (2021), we aimed to identify thalamic regions with a higher concentration of calbindin-rich matrix cells (supporting inter-network connectivity) and parvalbumin-rich core cells (supporting intra-network connectivity). Using the method of Müller *et al*. (2020), we leveraged MNI152 spatial maps of mRNA expression levels for PVALB and CALB1 proteins from the Allen Human Brain Atlas (Gryglewski *et al*., 2018). Voxel-wise mRNA levels were normalized and transformed into z-scores across all thalamic voxels. To differentiate between matrix and core cell-rich areas, a Difference z-score (D’ z-score = calbindin z-score − parvalbumin z-score) was calculated. D’ z-score > 0 indicated matrix cell-dominant voxels, while D’ z-score < 0 corresponded to core cell-rich regions. This D’ z-score map, displayed on an MNI152 template, was overlaid with voxel clusters associated with poorer performance on the FCSRT subtests. This map, only including voxels > 0 corresponding to calbindin-rich matrix cells was also overlaid with the overlap of all patient’s lesion divided into two groups: those with lesions that damaged the MTT and those with lesions that preserved it. The overlay also featured the manually segmented mask of the MTT on the MNI152 template.

### 2.7. Neurosynth

To assess whether our symptom mapping results aligned with previous literature, we used the Neurosynth query tool to identify memory-related voxels in human fMRI literature (Yarkoni *et al*., 2011). We queried all available memory-related keywords, including: “encoding memory,” “recall,” “retrieval,” “recognition memory,” “episodic memory” and “working memory.” To control for language effects, “language,” “verbal,” and “naming” keywords were also analyzed in the Supplementary Material. The Neurosynth tool generates voxel-wise maps based on a ‘uniformity test’, which reports z-scores (thresholded at z > 3) summarizing the likelihood of a voxel being reported in studies containing the selected keywords. This test performs a one-way ANOVA to determine whether activation at a given voxel occurs more frequently than expected under a uniform distribution across grey matter. To compare Neurosynth results with our symptom mapping findings, we overlapped the Neurosynth-derived maps with clusters of lesioned voxels identified in our analysis.

## 3. Results

### 3.1. Sociodemographic analyses

The two groups comprising 45 healthy subjects and 40 patients with isolated ischemic thalamic stroke were comparable in terms of gender (χ^2^ = 2.78, p = 0.10), age (t-test, p = 0.41) and years of education (t-test, p = 0.12) (Tab. 1). In addition, patients grouped by laterality of infarcts were comparable regarding lesion volume (Kruskal-Wallis: p = 0.285; L: 346 mm^3^, R: 405 mm^3^, B: 589 mm^3^). Overlaps of patients’ lesions by laterality of infarcts are illustrated in Supp. Fig. 1.

**Table 1:**
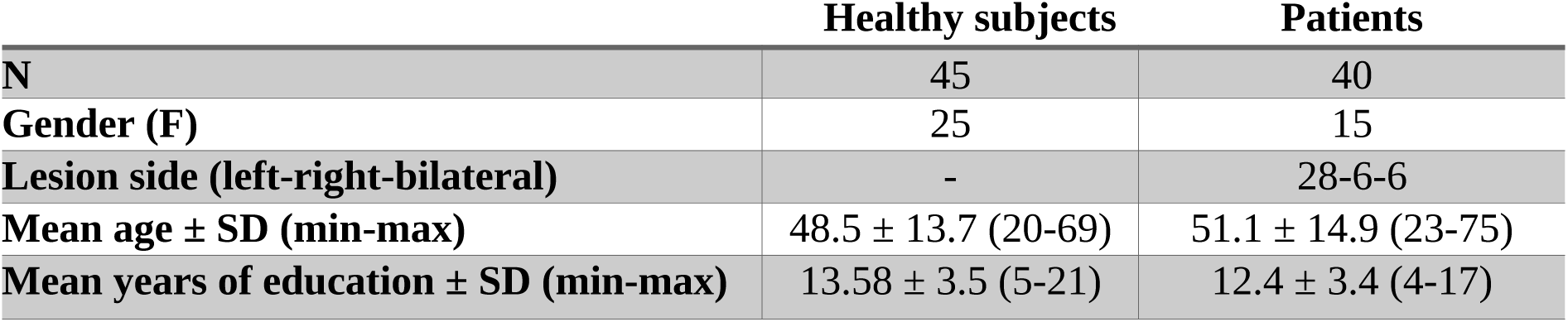
Sociodemographic information from the two groups of subjects.

|  | Healthy subjects | Patients |
| --- | --- | --- |
| **N** | 45 | 40 |
| **Gender (F)** | 25 | 15 |
| **Lesion side (left-right-bilateral)** | - | 28-6-6 |
| **Mean age ± SD (min-max)** | 48.5 ± 13.7 (20-69) | 51.1 ± 14.9 (23-75) |
| **Mean years of education ± SD (min-max)** | 13.58 ± 3.5 (5-21) | 12.4 ± 3.4 (4-17) |

### 3.2. Neuropsychological analyses and thalamic tract lesions

The primary goal of this analysis was to evaluate memory performance across groups, considering lesion laterality and MTT disruption. Significant group differences were observed in all subtests except Immediate Recall (see Fig. 2, Kruskal-Wallis followed by a Bonferroni-corrected Dunn’s test; Free Recall Sum: p < 0.001, η² = 0.38; Free Recall Total: p < 0.001, η² = 0.26; Delayed Free Recall: p < 0.001, η² = 0.39; Delayed Recall Total: p = 0.006, η² = 0.14).

**Figure 2:**
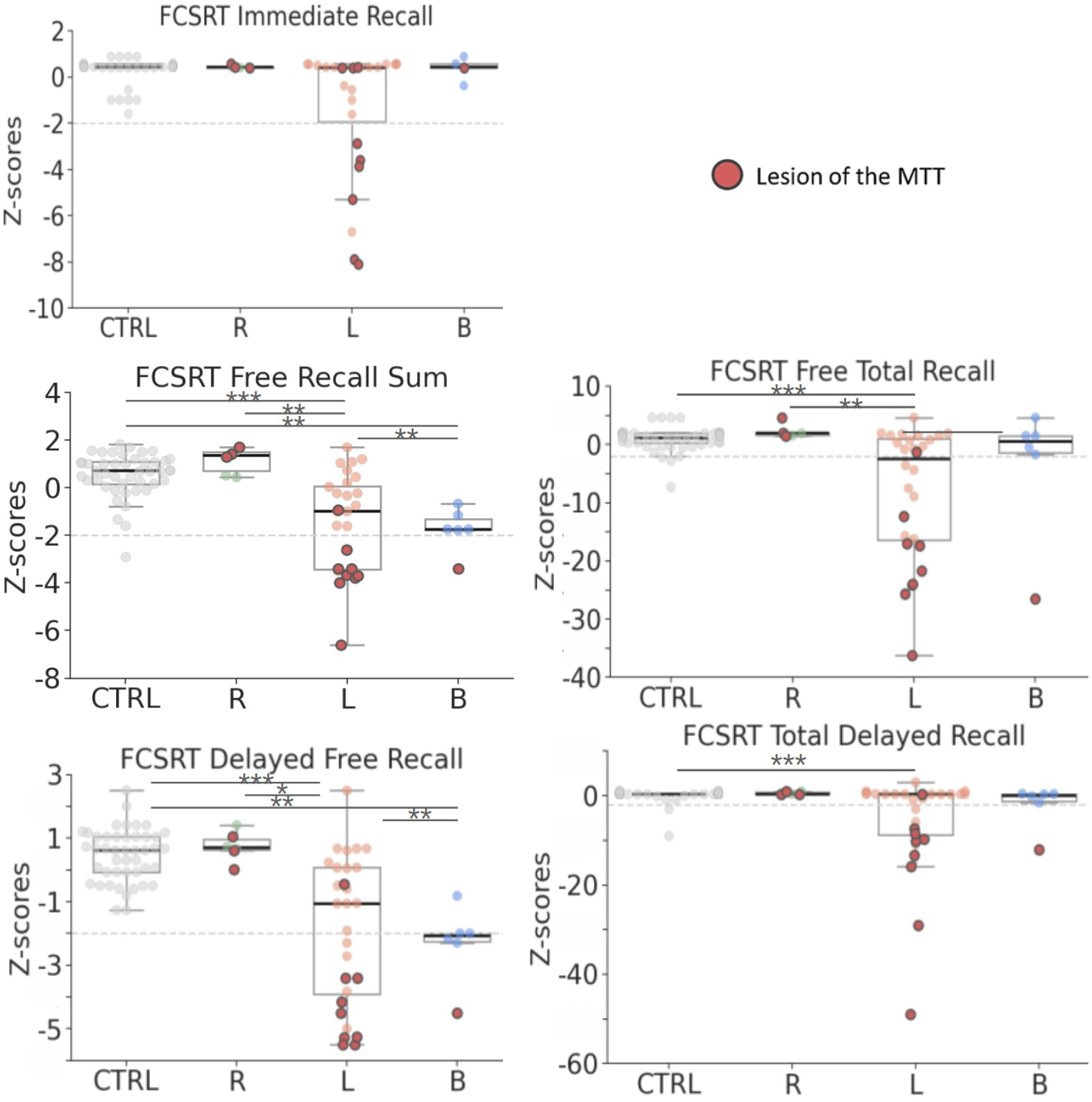
Boxplots displaying Z-scores for each FCSRT subtest by group and infarct laterality (Healthy subjects = 45; Patients: R = 6, L = 28, B = 6). The black line within each box indicates the median. Red circles represent patients with a lesion in the MTT (N = 13). The dashed gray line highlights a z-score of -2 SD, which is the usual threshold used in neuropsychological assessments. It is represented here for information only. Note that one data point is missing in the Immediate Recall subtest for the bilateral group. Statistical significance was assessed using Bonferroni-corrected Dunn’s test: *p < 0.05, **p < 0.01, ***p < 0.001.

Patients with left thalamic infarcts performed worse than healthy subjects across all subtests except Immediate Recall. They also performed worse than other patient groups in both the Free Recall Sum and Delayed Free Recall subtests. Bilateral lesions also led to deficits in these subtests. In contrast, patients with right infarcts were indistinguishable from healthy subjects.

13 patients had thalamic lesions extending into the MTT, 9 due to lesions of the left thalamus (mean lesion’s volume: 8 +- 4.8 mm^3^, min: 3, max: 19), 3 in the right group (mean lesion’s volume: 15 +- 16 mm^3^, min: 4, max: 33) and 1 with a bilateral thalamic lesion for which the MTT’s lesion was on the left side (lesion’s volume: 4 mm^3^). Patients with a left lesion of the MTT were the most impaired across all subtests, including Immediate Recall (Fig. 2). In addition, patients with left thalamic lesions and MTT disruptions (n = 9) were also significantly more impaired even when compared with patients with left thalamic lesions but preserved MTT (n = 19) (Mann-Whitney test: Immediate Recall, p=0.003, Rank-Biserial Correlation = 0.7; p<0.001 for all other subtests with a Rank-Biserial Correlation > 0.8).

Significant group differences were also observed across all tests for language performance (Supp. Fig. 2; Kruskal-Wallis: Literal Fluency, p = 0.003, η² = 0.155; Semantic Fluency, p < 0.001, η² = 0.28; Confrontation Naming, p = 0.001, η² = 0.18). Patients with left thalamic infarcts exhibited significantly lower language performance than healthy controls across all tests, while patients with bilateral infarcts showed impairment only in Semantic Fluency. Patients with right thalamic strokes were indistinguishable from healthy controls. However, unlike memory performance, MTT lesions were not consistently linked to language deficits. In addition, although group differences were observed, these deficits remained above -2 SD, except for one patient in both Fluencies and six patients in the Confrontation Naming tests (4/13 with a damaged MTT). This indicates that the language deficits were generally much less severe than the memory impairments.

### 3.3. Lesion symptom mapping

Symptom mapping analysis identifies lesioned voxel clusters associated with poorer performance in a specific test compared to patients with the same intact voxel. This technique was used to assess the effect of lesion location on memory performance for each FCSRT subtest among all patients, without regard to the laterality of infarcts. This analysis identified a cluster of lesioned voxels in the left thalamus significantly associated with poorer memory performance in the Free Recall Sum, Free Recall Total and Delayed Free Recall subtests (Fig. 3).

**Figure 3:**
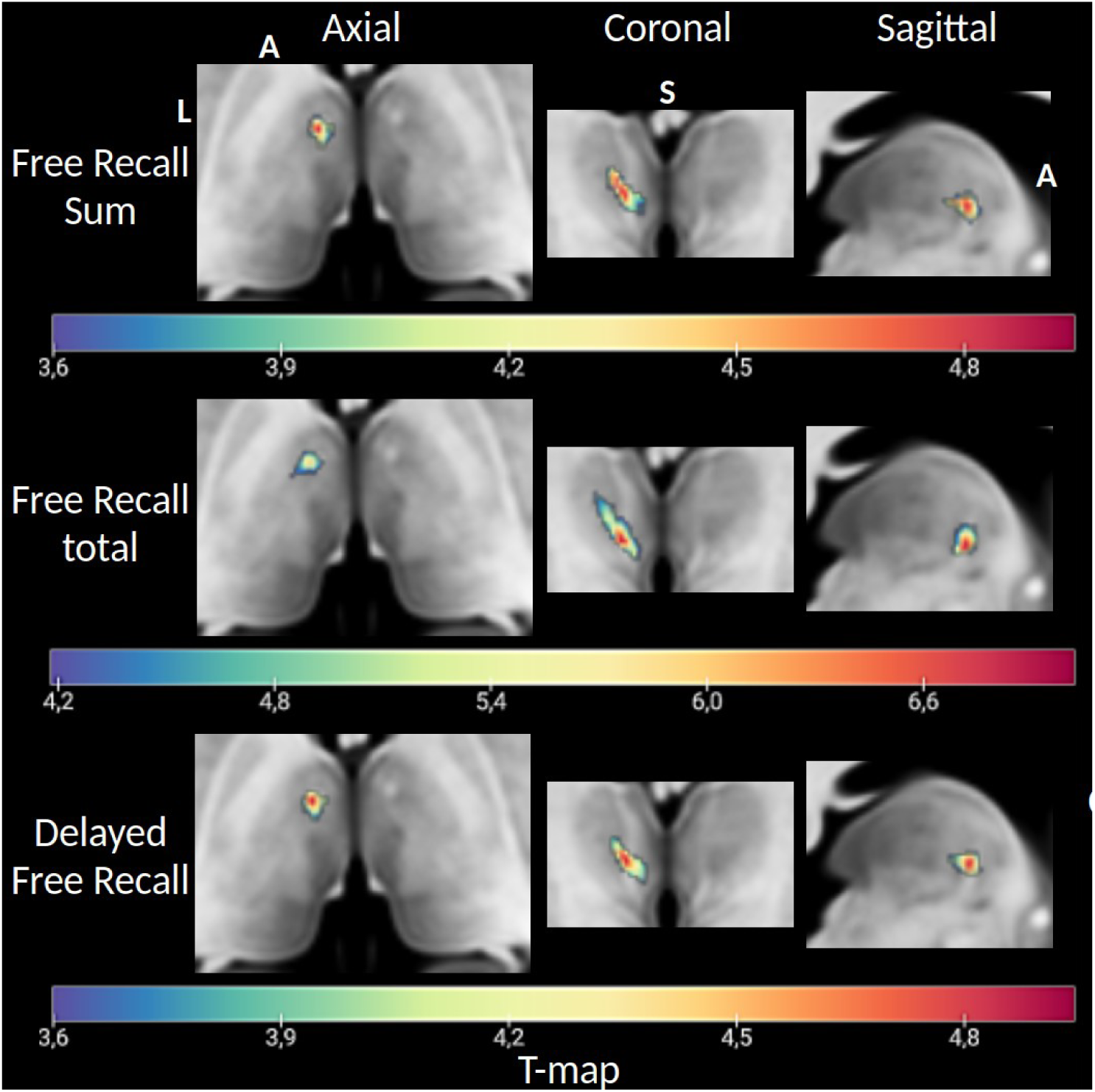
T-map of lesioned voxel clusters significantly associated with increased deficits in the FCSRT Free Recall Sum, Free Recall Total and Delayed Free Recall subtests on an MNI152 slice. L: Left, A: Anterior, S: Superior.

Given the consistency of the results between subtests, the cluster of voxels identified in the Delayed Free Recall was selected for visual representation on an MNI152 template slice and used to pinpoint lesion location using HIPS-THOMAS segmentation (Fig. 4). This lesioned voxel cluster was found to overlap with the ventral anterior (VA) and ventral lateral posterior (VLp) regions as well as with the MTT. Additionally, the voxels with the highest t-values, indicating the strongest association with memory deficits, almost perfectly overlapped with the MTT location.

**Figure 4:**
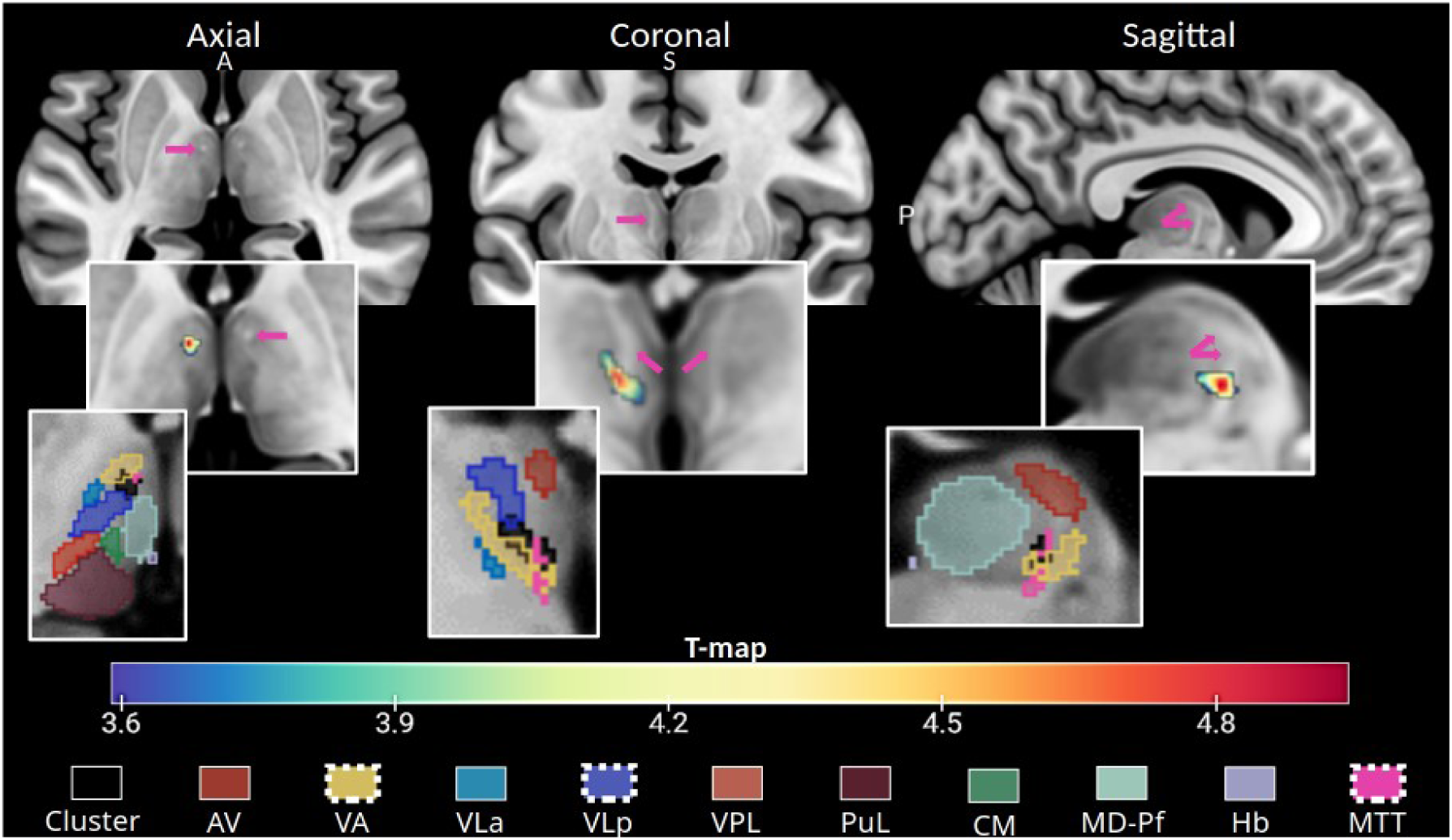
FCSRT cluster characterization. Visualization of the MTT along with the cluster of voxels associated with verbal memory deficits as measured by the FCSRT Delayed Free Recall. The upper section displays axial, coronal and sagittal MRI slices, with the MTT highlighted by pink arrows. Below, the same slices show an overlay of voxels significantly linked to increased deficits in the FCSRT Delayed Free Recall when lesioned. The lower section displays the same slices with an overlay of the HIPS-THOMAS segmentation of thalamic nuclei, and the cluster of voxels associated with verbal memory deficits in black (FCSRT Delayed Free Recall). Thalamic substructures containing voxels in this cluster include the VA (50%), VLp (<10%) and the MTT (<10%). AV: Anterior Ventral, VA: Ventral Anterior, VLa: Ventral Lateral anterior, VLp: Ventral Lateral posterior, MTT: mammillothalamic tract, A: Anterior, S: Superior, P: Posterior.

Notably, no significant voxel clusters were associated with language impairment in any of the language tasks (Supp. Fig. 3). In addition, the identified clusters of voxels, even without being significant, were not located in similar regions to those of the FCSRT and did not overlap with the MTT.

We ran a complementary independent disconnectome analysis to see if the lesioned voxel cluster significantly associated with increased deficits in the FCSRT (Free Recall sum) encompassed the MTT. We examined disrupted white matter tracts with the lesion mask as a seed using tractography in 10 healthy subjects to generate a map of interrupted bundles passing through the lesion (see Methods). Results are represented as a t-map (Fig. 5A) and binarized with the MTT segmentation overlap for location comparison (Fig. 5B). As expected, it visually overlapped with the MTT location. In addition, the cluster of disconnected voxels was associated with left anterior thalamic radiation disruption (100% probability) and left frontostriatal tract disruption (52%).

**Figure 5:**
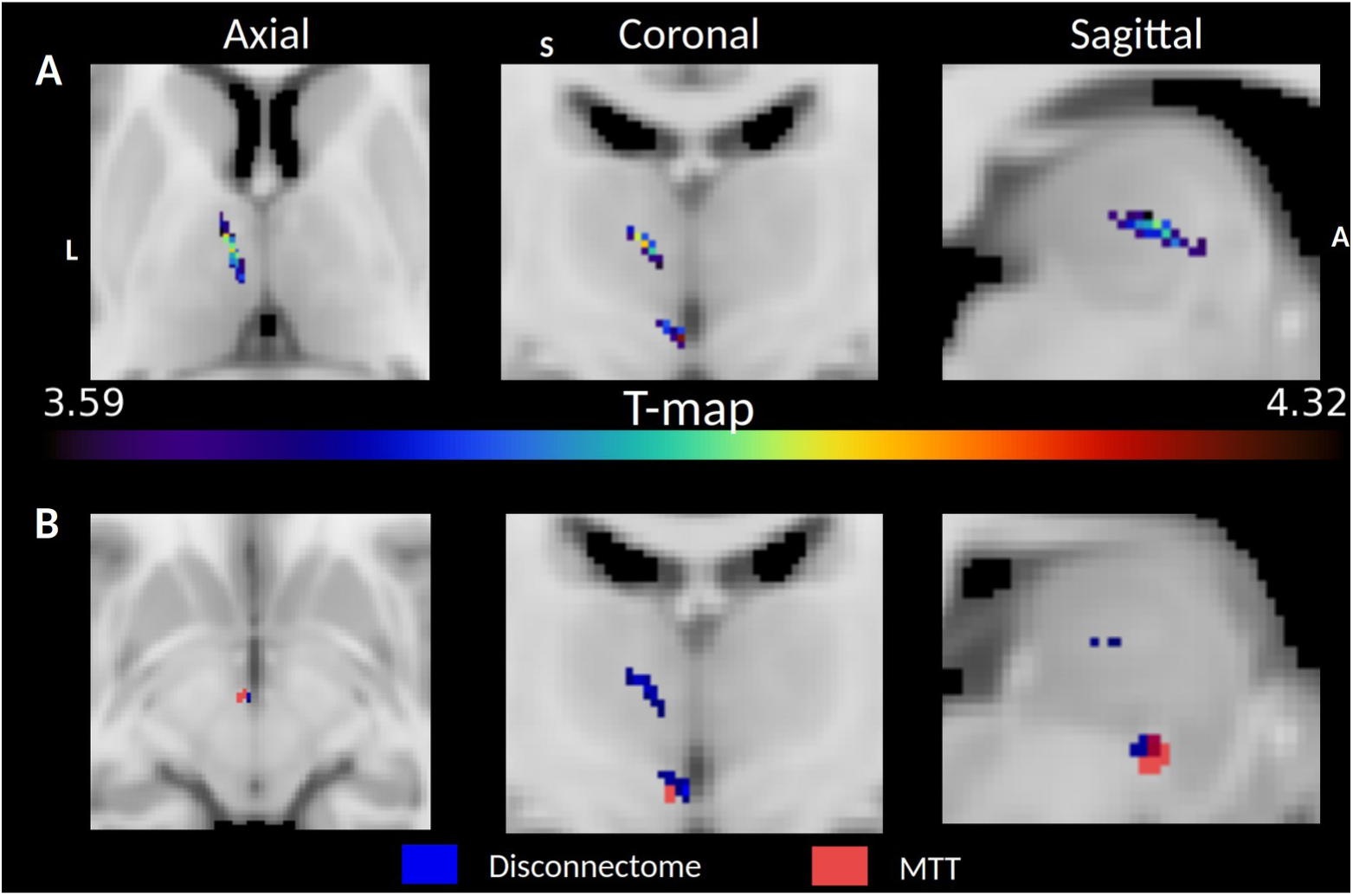
A- T-map of disconnected voxels associated with significantly poorer FCSRT Delayed Free Recall performances on an MNI152 slice. B- Same T-map, which was binarized and overlapped with the MTT HIPS-THOMAS segmentation of the MNI152 template. Both locations almost perfectly overlap, suggesting a disconnection of the MTT.

### 3.4. Testing another hypothesis

Some authors have proposed that the calbindin-rich anterior thalamus might serve as a critical hub for higher cognitive functions, suggesting that, if this region is lesioned, the patients would show a broad range of deficits. Here, we investigated whether this, rather than lesions to the MTT itself, could account for the patients’ memory impairment observed in patients with left thalamic stroke. Reproducing the findings of Hwang *et al*. (2021), we identified thalamic areas with greater relative concentrations of parvalbumin (core cells, high intra-network connectivity, provincial hub properties) or calbindin (matrix cells, high internetwork connectivity, connector hub properties). We then overlaid on these areas the cluster of voxels linked to FCSRT deficits (Fig. 6A), showing an overlap with calbindin-rich regions. Next, we overlaid the structural lesions mask of the group of patients with either damaged, or preserved, left MTT (Fig. 6B). Both sets of lesions overlapped with the calbindin-rich areas. As the patients with left lesioned MTT showed much more severe memory impairment, these results indicate that impaired memory observed in patients with damage to the MTT can only result from disruption of the MTT itself and not from lesions to the hub.

**Figure 6:**
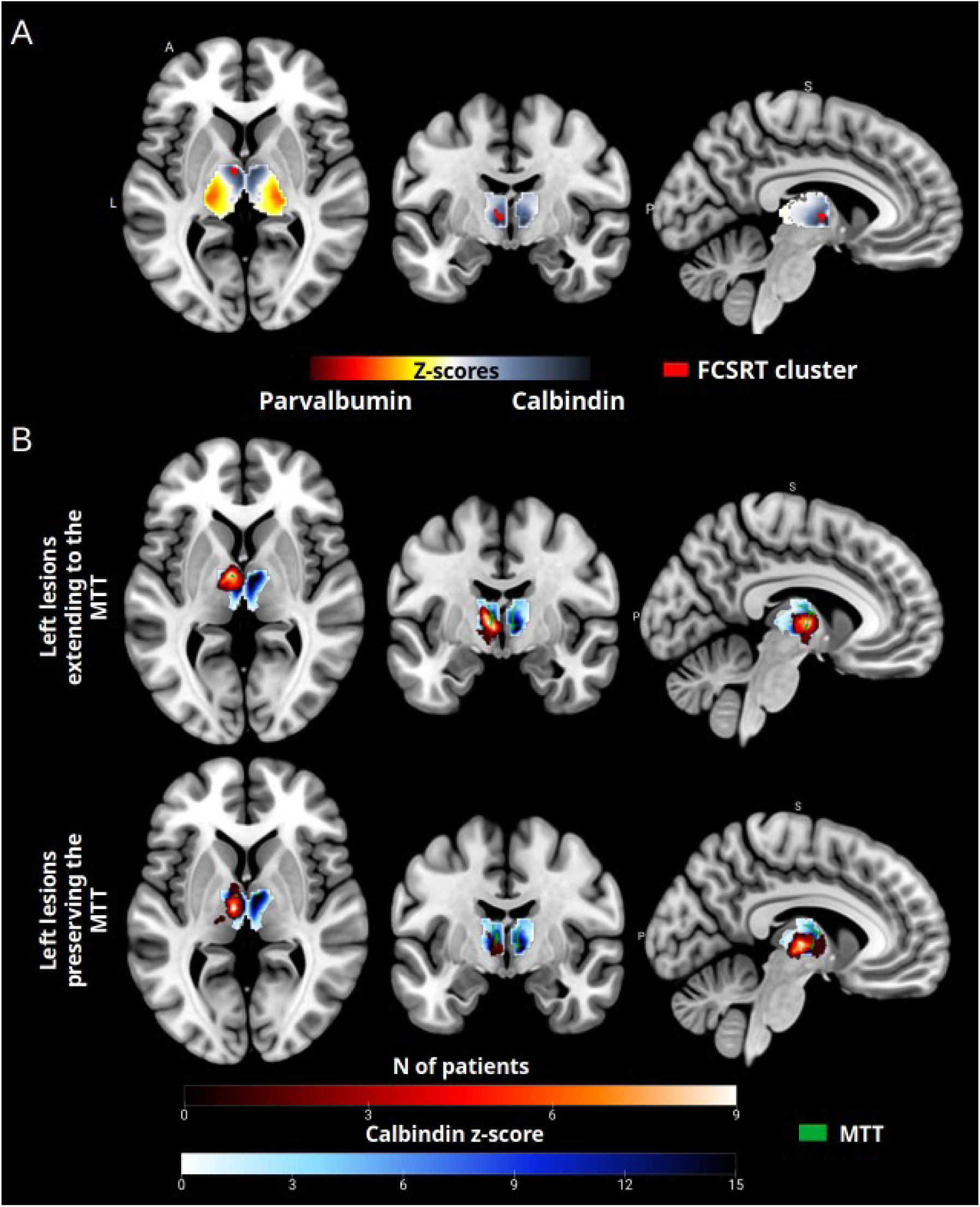
A- Relative expression of parvalbumin and calbindin in the thalamus (D’ z-score) overlaid with voxels associated with greater deficits in the FCSRT test (cf Fig. 4) in cases of thalamic stroke (FCSRT cluster). B- Relative expression of calbindin in the thalamus (D’ z-score > 0), overlaid with patient lesions. The upper figure shows lesions that extend into the left MTT, while the lower figure shows lesions that preserve the left MTT. The manual segmentation of the MTT is also included. A: Anterior, P: Posterior, R: Right, S: Superior.

### 3.5. Generalizability of the results

The Neurosynth query tool identifies voxels associated with specific keywords from the human fMRI literature. To validate and test the generalizability of our findings, we compared the identified FCSRT voxel cluster with Neurosynth-generated meta-analytic maps of memory-related keywords.

This cluster overlapped with all memory-related keywords except “episodic memory”, with the highest concordance observed in coronal slices for “recall” and “retrieval”, covering 90% of the cluster. Keyword “recognition” and “encoding” encompassed 67%, while “working memory” accounted for 44% (Fig. 7). To control for language effects, related keywords are analyzed and presented in Supp Fig. 4. An overlap was observed for “language” (67%), while no correspondence was found for “verbal” (22%) or “naming” (0%) keywords.

**Figure 7:**
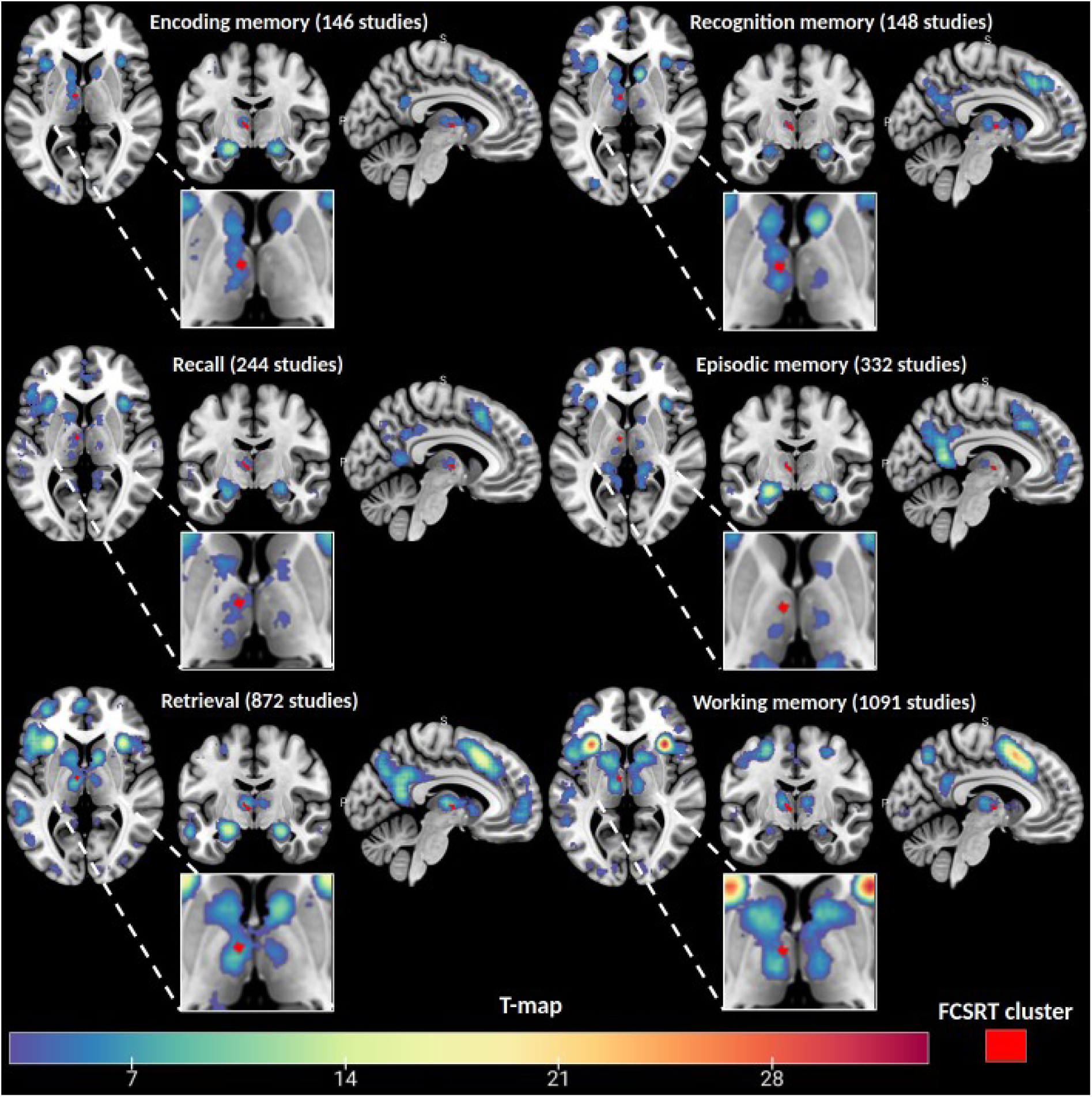
Visual representation of results from the Neurosynth query tool, displayed as T-maps of significant voxels associated with memory-related keywords from fMRI human studies. The cluster of voxels associated with poorer performance on the FCSRT Delayed Free Recall subtest is overlaid in red for comparison.

## 4. Discussion

This study provides robust evidence that the MTT plays a pivotal role in memory impairments following thalamic strokes. It challenges the traditional focus on thalamic nuclei and the recently proposed hub view of the thalamus to explain memory deficits.

We assessed whether the MTT was damaged on each individual MRI for a group of patients with isolated thalamic stroke. Symptom mapping analysis revealed that memory impairments were associated with lesions overlapping the ventral anterior and ventral lateral posterior nuclei. However, the voxels most strongly correlated with memory deficits were predominantly located at the MTT location. This finding was further supported by a disconnectome analysis, underlining the importance of the MTT in thalamic stroke outcomes. In addition, we tested whether memory impairment could be related to lesions in the anterior calbindin-rich area of the thalamus. The results indicated that these deficits could only be attributed to damage to the MTT itself, rather than disruption of a connector hub. Finally, our findings, analyzed using a single well-known verbal memory test, appear to generalize to other memory tests, as supported by meta-analytic maps of memory-related keywords (Neurosynth query).

### Laterality of Thalamic Stroke and Verbal Memory

FCSRT performance analysis, based on infarct laterality, revealed that patients with left-sided thalamic strokes exhibited significant impairments in a free and cued recall task compared to healthy controls. Impairments were particularly pronounced in patients with left MTT damage. These findings suggest that memory deficits in these patients stem from ineffective encoding or storage, resulting in weak or absent memory traces that cannot be retrieved even with cues. The most pronounced impairments were observed in Free Recall Sum but also on the Delayed (20 min) Free Recall subtest. The FCSRT is a relatively easy test for healthy subjects, with a ceiling effect as, by the time of the delayed recall, each word has been recalled or rehearsed four times. This indicates that patients failing on the FCSRT show significant verbal memory deficits.

In contrast, patients with right thalamic infarcts performed similarly to healthy controls, highlighting, as expected, a strong hemispheric asymmetry in verbal memory function. This implies that the left thalamus plays a more critical role in verbal processing, probably due to stronger connections with language-related cortical areas (Crosson, 1985). Bilateral thalamic lesions, although less frequent, also led to deficits across multiple FCSRT subtests.

Patients with left-sided strokes also presented language deficits, albeit to a lesser extent than memory impairments, mainly affecting one patient when observing individual performances. This aligns with previous neuropsychological studies observing language deficits in some patients and semantic deficits in case of left thalamic lesions (Nadeau & Crosson, 1997; De Witte *et al*., 2011; Crosson, 2013; Pergola *et al*., 2013).

### Thalamic Tracts, Verbal Memory and the FCSRT

The MTT connects the hippocampus to the anterior thalamic nucleus through the mammillary bodies, playing a crucial role in episodic memory. Our findings confirm that, in all memory subtests, the poorest memory performances were observed in patients with MTT disruptions, emphasizing the MTT’s essential role in memory encoding, retrieval and consolidation. These results align with a review of 35 studies (60 patients with thalamic infarcts), demonstrating that MTT damage is strongly linked to memory impairment (Van der Werf *et al*., 2000). Furthermore, the laterality of the MTT’s lesion was a more significant predictor of memory deficits than its lesion volume. Indeed, no deficit was observed in patients with right-sided lesions, even though this group included the patient with the largest MTT lesion volume (33 mm³). This phenomenon can be explained by the small size of the MTT, where even a lesion as small as 1 mm³ (1 voxel) can potentially interrupt its connectivity and lead to memory deficits.

### Thalamic Area Associated with Verbal Memory

One of the most compelling findings of this study is the identification of a specific cluster of lesioned voxels associated with verbal memory deficits. This region remained consistent across the Free Recall Sum, Free Recall Total and Delayed Free Recall subtests, although it did not reach significance for language tests. This lesion cluster was primarily located in the left anterior-lateral thalamus, overlapping with the ventral anterior and ventral lateral posterior nuclei, and especially the MTT. The identified hub aligns closely with previous research, which highlights the anterior lateral and medial thalamic regions as key integrators in memory, executive function and language networks (Van der Werf *et al*., 2003; Scharf *et al*., 2020; Hwang *et al*., 2021; Vidal *et al*., 2025).

In addition, the ventral anterior and ventral lateral posterior nuclei may also contribute to lexical and semantic processing, potentially explaining impairments in Confrontation Naming and Fluency tests in a subset of patients with lesions in these regions (Nishio *et al*., 2014). The Selective Engagement Model (Nadeau & Crosson, 1997) posits that the ventral anterior nucleus plays a role in word selection (Crosson *et al*., 2007), linking semantic and verbal memory deficits. However, the symptom mapping analyses did not yield significant results for language tests. This suggests that the thalamic regions affected by lesions in our dataset may not be specifically involved in language processing. This finding further implies that the verbal memory deficits associated with these thalamic lesions cannot be solely attributed to impairments in language processing.

### Thalamic Connector Hub and Provincial Hub

The voxel cluster associated with verbal memory deficits overlapped with a calbindin-rich thalamic region. However, lesions within these connector hubs alone do not fully explain memory impairments since patients with a preserved MTT but with lesions in the same connector hub showed no significant memory impairments. This suggests that MTT disruption, rather than damage to the hub, was the primary driver of memory impairments. While connector hubs are essential for network integration, white matter tracts like the MTT are probably the key contributors to memory deficits following thalamic strokes.

Given the MTT’s critical role in memory processing, determining whether this tract is affected in thalamic stroke is essential not only to understand the origins of post-stroke memory deficits but also for clinical purposes. Specifically, identifying MTT involvement can help stratify patients at higher risk of memory impairment, and guide more targeted therapeutic and rehabilitative interventions.

### Disruption of White Matter Tracts and Cognitive Deficits

Given its significant role, identifying MTT lesions is essential for understanding cognitive impairments associated with thalamic damage. This raises a broader question: Should other thalamic tracts be considered following thalamic strokes? Two less-studied thalamic tracts may be of importance. The ventral amygdalo-fugal pathway, located lateral to the MTT, connects the amygdala to the mediodorsal nucleus of the thalamus, a key component of the limbic system (Graff-Radford *et al*., 1990; Erkan *et al*., 2024). Meanwhile, the interthalamic adhesion (IA) links the two thalami across the third ventricle in approximately 80% of the population (Borghei *et al*., 2021; Wong *et al*., 2021; Vidal *et al*., 2024b) and may play a compensatory role in preserving cognitive function, particularly in verbal memory, following thalamic damage (Tresniak *et al*., 2016; Vidal *et al*., 2024b). To fully understand the thalamus’s contribution to cognition, future studies should assess the integrity of these pathways.

### Limitations

Despite including a relatively large cohort of patients with isolated thalamic strokes, the rarity of such cases, particularly with isolated bilateral thalamic strokes (Percheron, 1973), may limit the generalizability of our findings. The lesioned nuclei in our cohort were predominantly supplied by paramedian arteries. This overrepresentation is partly attributable to our inclusion criteria. Among thalamic vascular territories, the paramedian territory, which includes the MTT, is the second most frequently affected, representing around 35% of all thalamic strokes, after the inferolateral territory (45%) and preceding the anterior (12%) and posterior (8%) territories (Schmahmann *et al*., 2003). Expanding research to include a broader range of impaired arterial territories could provide a more comprehensive understanding of thalamic stroke impact on memory. Anatomical MRI also cannot fully delineate tract pathways or quantify tract disruption. Diffusion-weighted imaging and advanced tractography methods (e.g., probabilistic fiber tracking) would provide greater accuracy. However, improved spatial resolution is needed to assess the MTT integrity.

## 5. Conclusion

This study underlines the critical role of the MTT in verbal memory impairments following left thalamic strokes. Our findings challenge the prevailing focus on thalamic nuclei and hubs over individual white matter tracts in studying thalamic function. While connector hubs enriched with matrix cells are pivotal for inter-network communication, our results indicate that damage to the MTT is a stronger predictor of verbal memory impairments. These findings emphasize the need to consider white matter tracts running through the thalamus when studying the impact of pathology on the thalamus.

## 6. Data availability

Datasets are available from the corresponding author upon reasonable request.

## 7. Authors contributions

J.P.V. and E.J.B. designed the study. J.P.V. performed the analyses. J.P.V wrote the manuscript with the support of E.J.B. J.P, J.F.A, P.P., L.D, participated in recruitment, in acquisition of neuropsychological and neuroimaging data, and in the revision of the manuscript.

## 8. Acknowledgements

We would like to acknowledge the contribution of Michel-Thiebaut de Schotten and Chris Foulon for their valuable assistance with the use of the BCBtoolkit software and FSL processing.

## 9. Competing Interest

The authors have no conflicts of interest relevant to this manuscript to disclose.

## Supplementary Figure

**Supplementary figure 1:**
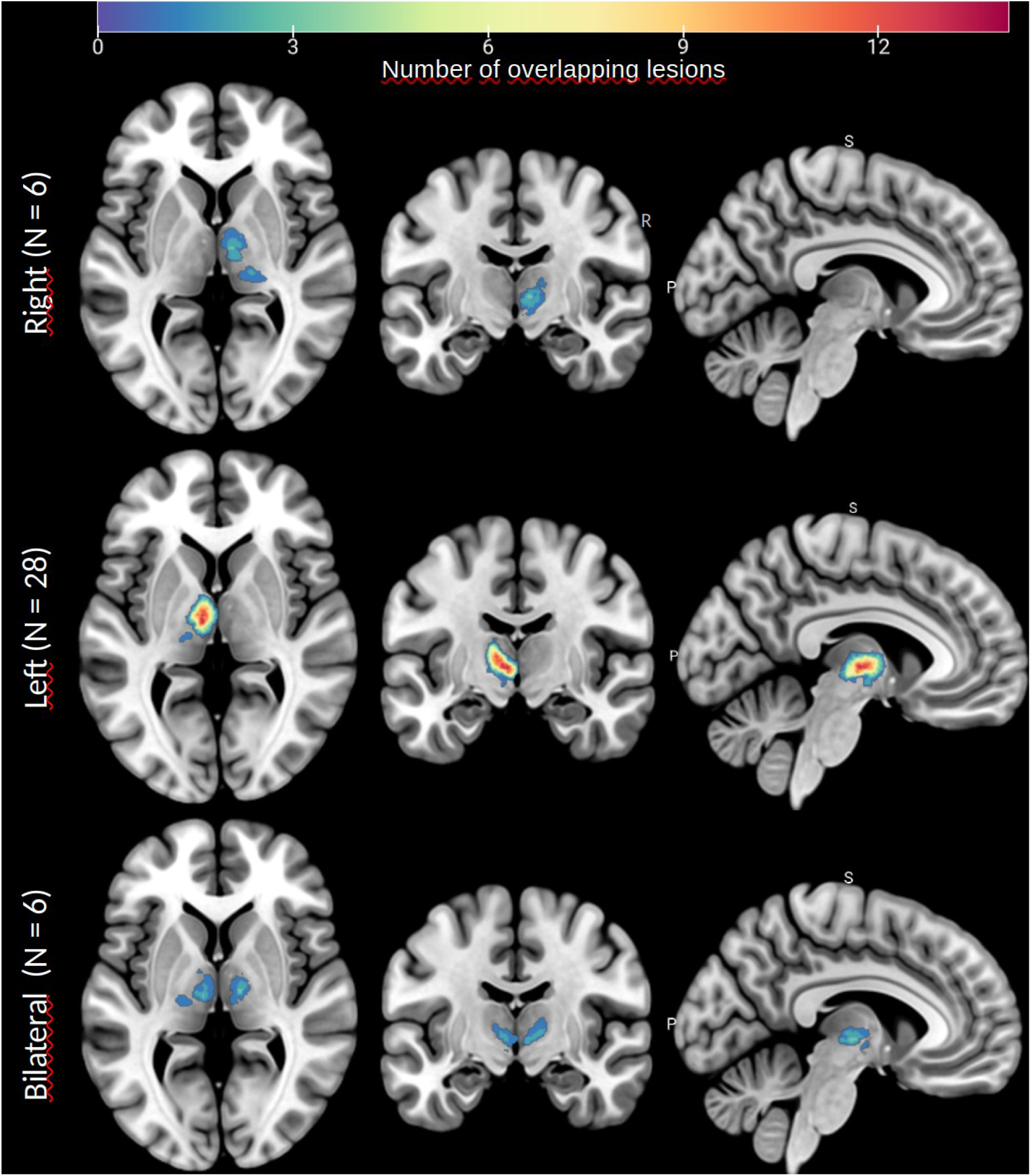
Representation of all lesions from the 40 patients on the MNI152 template after normalization and by infarct laterality. R: Right; S: Superior; P: Posterior

**Supplementary figure 2:**
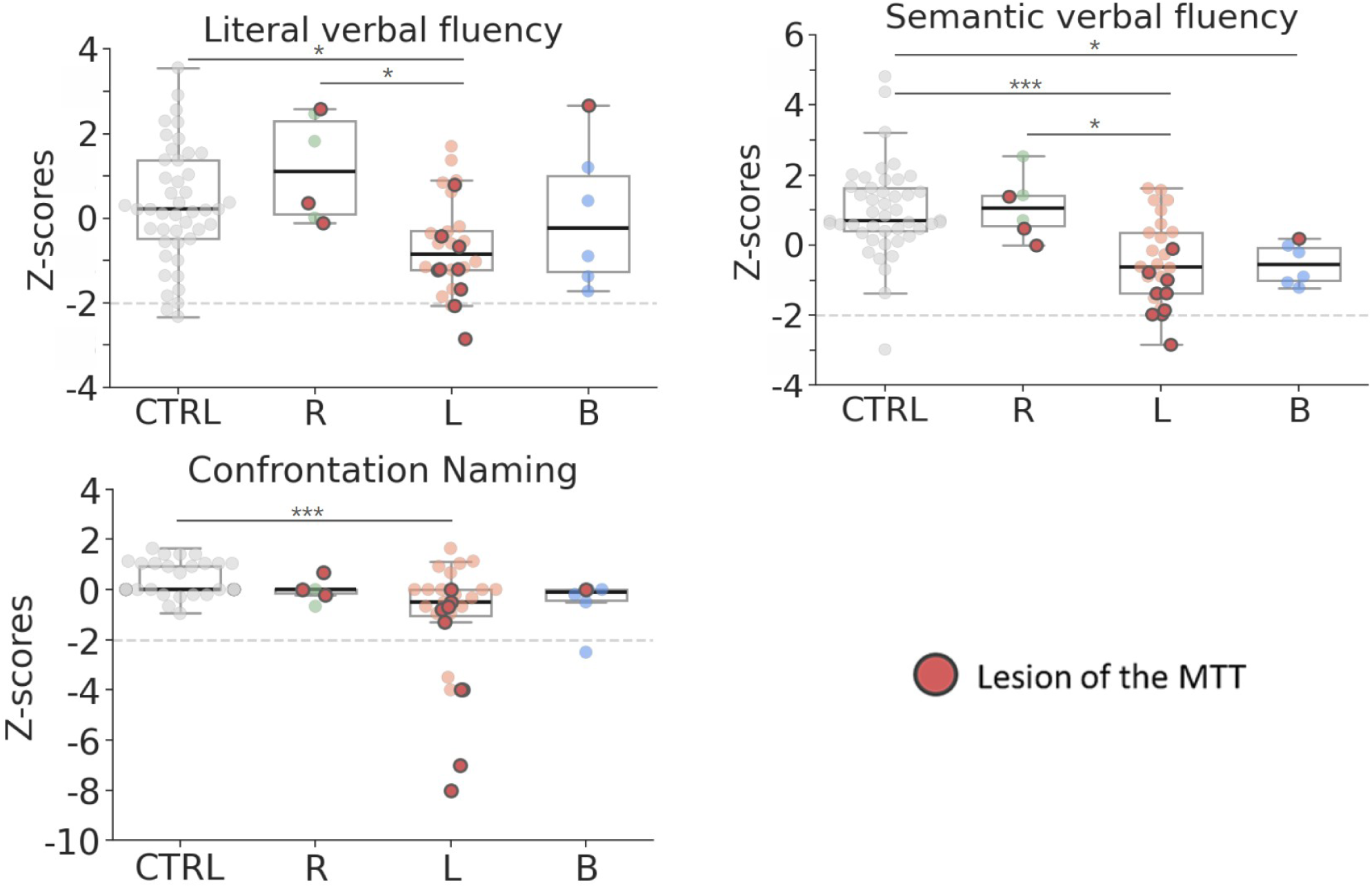
Boxplots displaying Z-scores for literal and semantic verbal fluency, and for the confrontation naming test, by group and infarct laterality (healthy subjects = 45; Right = 6, Left = 28, Bilateral = 6). The black line within each box indicates the median. The gray dashed line highlights a z-score of -2 SD usually thresholding significant deficits. Red rounds represent patients with a lesion in the MTT. Statistical significance was assessed using Bonferroni-corrected Dunn’s test: *p < 0.05, **p < 0.01, ***p < 0.001.

**Supplementary figure 3:**
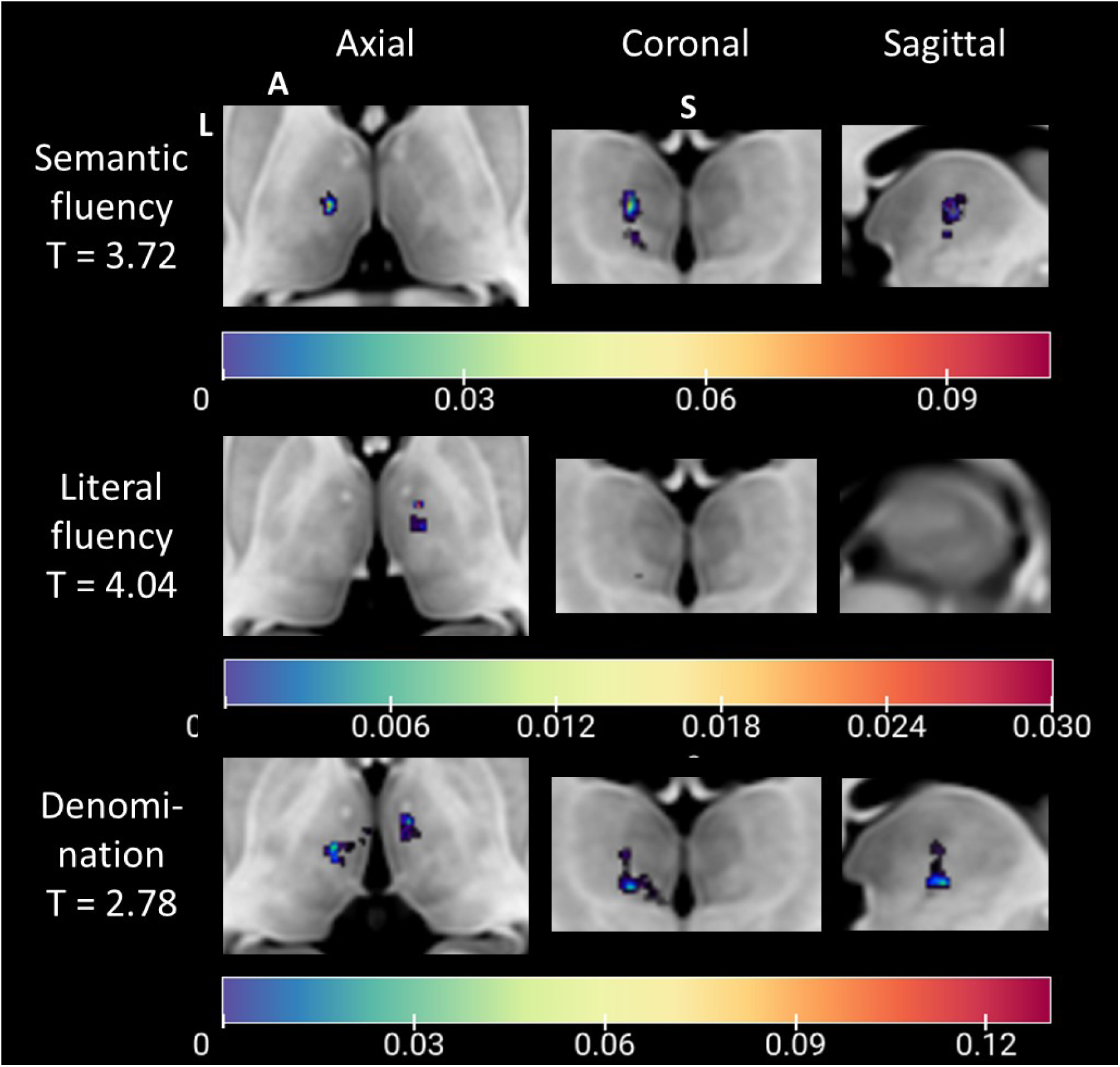
T-map of lesioned voxel clusters associated with deficits in Semantic Fluency, Literal Fluency or Confrontation Naming tests overlaid on an MNI152 slice. The T-value (T) represents the threshold for significance, which was not exceeded in any of these tests as reflected by the scales. L: Left, A: Anterior, S: Superior.

**Supplementary figure 4:**
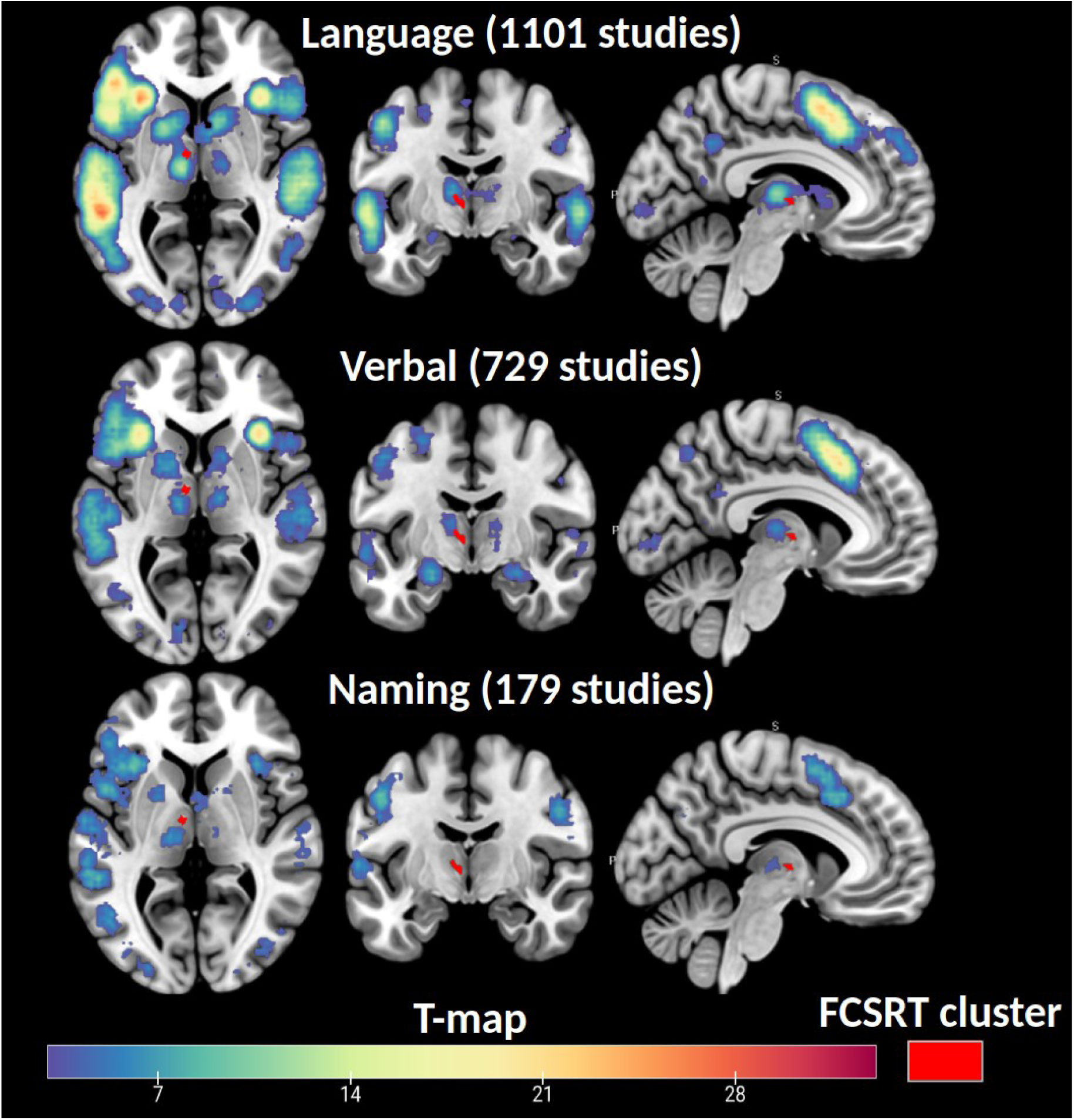
Visual representation of results from the Neurosynth query tool, displayed as T-maps of significant voxels associated with memory-related keywords from fMRI human studies. The cluster of voxels associated with poorer performance on the FCSRT Delayed Free Recall subtest is overlaid in red for comparison.

## Notes

### Competing Interest Statement

The authors have declared no competing interest.

## References

Aggleton, J. P., & Brown, M. W. (1999). Episodic memory, amnesia, and the hippocampal– anterior thalamic axis. Behavioral and brain sciences, 22(3), 425–444.

Aggleton, J. P., & Brown, M. W. (2006). Interleaving brain systems for episodic and recognition memory. Trends in cognitive sciences, 10(10), 455–463.

Aggleton, J. P., Vann, S. D., & O’Mara, S. M. (2023). Converging diencephalic and hippocampal supports for episodic memory. Neuropsychologia, 191, 108728.

Bachy-Langedock, N. (1989). Batterie d’examen des troubles de la dénomination (ExaDé). Brussels: Editions du Centre de Psychologie Appliquée.

Borges, A., Piracha, A., Sani, S. (2021). Prevalence and anatomical characteristics of the human massa intermedia. Brain Structure and Function, 226, 471–480.

Carlesimo, G. A., Serra, L., Fadda, L., Cherubini, A., Bozzali, M., & Caltagirone, C. (2007). Bilateral damage to the mammillo-thalamic tract impairs recollection but not familiarity in the recognition process: a single case investigation. Neuropsychologia, 45(11), 2467–2479.

Cipolotti, L., Husain, M., Crinion, J., Bird, C. M., Khan, S. S., Losseff, N., … & Leff, A. P. (2008). The role of the thalamus in amnesia: a tractography, high-resolution MRI and neuropsychological study. Neuropsychologia, 46(11), 2745–2758.

Crosson, B. (1985). Subcortical functions in language: a working model. Brain and language, 25(2), 257–292.

Crosson, B. (2013). Thalamic mechanisms in language: a reconsideration based on recent findings and concepts. Brain and language, 126(1), 73–88.

Crosson, B., Fabrizio, K. S., Singletary, F., Cato, M. A., Wierenga, C. E., Parkinson, R. B., … & Rothi, L. J. G. (2007). Treatment of naming in nonfluent aphasia through manipulation of intention and attention: A phase 1 comparison of two novel treatments. Journal of the International Neuropsychological Society, 13(4), 582–594.

Danet, L., Barbeau, E. J., Eustache, P., Planton, M., Raposo, N., Sibon, I., Albucher, J. F., Bonneville, F., Peran, P., & Pariente, J. (2015). Thalamic amnesia after infarct. Neurology, 85(24), 2107–2115.

de Jong, L. W., van der Hiele, K., Veer, I. M., Houwing, J. J., Westendorp, R. G. J., Bollen, E. L. E. M., … & van der Grond, J. (2008). Strongly reduced volumes of putamen and thalamus in Alzheimer’s disease: an MRI study. Brain, 131(12), 3277–3285.

De Witte, L., Brouns, R., Kavadias, D., Engelborghs, S., De Deyn, P. P., & Mariën, P. (2011). Cognitive, affective and behavioural disturbances following vascular thalamic lesions: a review. Cortex, 47(3), 273–319.

Dillingham, C. M., Frizzati, A., Nelson, A. J., & Vann, S. D. (2015). How do mammillary body inputs contribute to anterior thalamic function?. Neuroscience & Biobehavioral Reviews, 54, 108–119.

Dillingham, C. M., Milczarek, M. M., Perry, J. C., & Vann, S. D. (2021). Time to put the mammillothalamic pathway into context. Neuroscience & Biobehavioral Reviews, 121, 60–74.

Erkan, B., Hergünsel, B., Barut, O., Saygı, T., Kocak, B., Güngör, A., … & Tanriover, N. (2024). Ventral amygdalofugal pathway as an integrated surgically important network: microsurgical anatomy and segmentation based on fiber dissection. Journal of Neurosurgery, 1(aop), 1–15.

Foulon, C., Cerliani, L., Kinkingnehun, S., Levy, R., Rosso, C., Urbanski, M., … & Thiebaut de Schotten, M. (2018). Advanced lesion symptom mapping analyses and implementation as BCBtoolkit. Gigascience, 7(3), giy004.

Forkel, S. J., de Schotten, M. T., Kawadler, J. M., Dell’Acqua, F., Danek, A., & Catani, M. (2014). The anatomy of fronto-occipital connections from early blunt dissections to contemporary tractography. Cortex, 56, 73–84.

Godefroy, O. GREFEX. (2008). Fonctions exécutives et pathologies neurologiques et psychiatriques.

Gratton, C., Nomura, E. M., Pérez, F., & D’Esposito, M. (2012). Focal brain lesions to critical locations cause widespread disruption of the modular organization of the brain. Journal of cognitive neuroscience, 24(6), 1275–1285.

Grober, E., Buschke, H. C. H. B. S., Crystal, H., Bang, S., & Dresner, R. (1988). Screening for dementia by memory testing. Neurology, 38(6), 900–900.

Grober, E., Lipton, R. B., Hall, C., & Crystal, H. (2000). Memory impairment on free and cued selective reminding predicts dementia. Neurology, 54(4), 827–832.

Graff-Radford, N. R., Tranel, D., Van Hoesen, G. W., & Brandt, J. P. (1990). Diencephalic amnesia. Brain, 113(1), 1–25.

Gryglewski, G., Seiger, R., James, G. M., Godbersen, G. M., Komorowski, A., Unterholzner, J., … & Lanzenberger, R. (2018). Spatial analysis and high-resolution mapping of the human whole-brain transcriptome for integrative analysis in neuroimaging. Neuroimage, 176, 259–267.

Hwang, K., Bertolero, M. A., Liu, W. B., & D’Esposito, M. (2017). The human thalamus is an integrative hub for functional brain networks. Journal of Neuroscience, 37(23), 5594–5607.

Hwang, K., Shine, J. M., Bruss, J., Tranel, D., & Boes, A. (2021). Neuropsychological evidence of multi-domain network hubs in the human thalamus. Elife, 10, e69480.

Jones, E. G. (2001). The thalamic matrix and thalamocortical synchrony. Trends in neurosciences, 24(10), 595–601.

Jones, E. G. (2009). The origins of cortical interneurons: mouse versus monkey and human. Cerebral cortex, 19(9), 1953–1956.

Krauth, A., Blanc, R., Poveda, A., Jeanmonod, D., Morel, A., & Székely, G. (2010). A mean three-dimensional atlas of the human thalamus: generation from multiple histological data. Neuroimage, 49(3), 2053–2062.

Kwon, H. G., Hong, J. H., & Jang, S. H. (2010). Mammillothalamic tract in human brain: diffusion tensor tractography study. Neuroscience letters, 481(1), 51–53.

Low, A., Mak, E., Malpetti, M., Chouliaras, L., Nicastro, N., Su, L., … & O’Brien, J. T. (2019). Asymmetrical atrophy of thalamic subnuclei in Alzheimer’s disease and amyloid-positive mild cognitive impairment is associated with key clinical features. Alzheimer’s & Dementia: Diagnosis, Assessment & Disease Monitoring, 11, 690–699.

Müller, E. J., Munn, B., Hearne, L. J., Smith, J. B., Fulcher, B., Arnatkevičiūtė, A., … & Shine, J. M. (2020). Core and matrix thalamic sub-populations relate to spatio-temporal cortical connectivity gradients. NeuroImage, 222, 117224.

Morel, A. (2007). Stereotactic atlas of the human thalamus and basal ganglia.

Nadeau, S. E., & Crosson, B. (1997). Subcortical aphasia. Brain and language, 58(3), 355–402.

Nishio, Y., Hashimoto, M., Ishii, K., Ito, D., Mugikura, S., Takahashi, S., & Mori, E. (2014). Multiple thalamo-cortical disconnections in anterior thalamic infarction: implications for thalamic mechanisms of memory and language. Neuropsychologia, 53, 264–273.

Pergola, G., Bellebaum, C., Gehlhaar, B., Koch, B., Schwarz, M., Daum, I., & Suchan, B. (2013). The involvement of the thalamus in semantic retrieval: a clinical group study. Journal of Cognitive Neuroscience, 25(6), 872–886.

Percheron, G. (1973). The anatomy of the arterial supply of the human thalamus and its use for the interpretation of the thalamic vascular pathology. Zeitschrift für Neurologie, 205, 1–13.

Rorden, C., & Brett, M. (2000). Stereotaxic display of brain lesions. Behavioural neurology, 12(4), 191–200.

Saygin, Z. M., Kliemann, D., Iglesias, J. E., van der Kouwe, A. J., Boyd, E., Reuter, M., … & Alzheimer’s Disease Neuroimaging Initiative. (2017). High-resolution magnetic resonance imaging reveals nuclei of the human amygdala: manual segmentation to automatic atlas. Neuroimage, 155, 370–382.

Scharf, A. C., Gronewold, J., Todica, O., Moenninghoff, C., Doeppner, T. R., de Haan, B., … & Hermann, D. M. (2022). Evolution of neuropsychological deficits in first-ever isolated ischemic thalamic stroke and their association with stroke topography: a case-control study. Stroke, 53(6), 1904–1914.

Schmahmann, J. D. (2003). Vascular syndromes of the thalamus. Stroke, 34(9), 2264–2278.

Segobin, S., Haast, R. A., Kumar, V. J., Lella, A., Alkemade, A., Bach Cuadra, M., … & Hornberger, M. (2024). A roadmap towards standardized neuroimaging approaches for human thalamic nuclei. Nature Reviews Neuroscience, 1–17.

Sherman, S. M. (2016). Thalamus plays a central role in ongoing cortical functioning. Nature neuroscience, 19(4), 533–541.

Souter, N. E., Wang, X., Thompson, H., Krieger-Redwood, K., Halai, A. D., Lambon Ralph, M. A., … & Jefferies, E. (2022). Mapping lesion, structural disconnection, and functional disconnection to symptoms in semantic aphasia. Brain Structure and Function, 227(9), 3043–3061.

Su, J. H., Thomas, F. T., Kasoff, W. S., Tourdias, T., Choi, E. Y., Rutt, B. K., & Saranathan, M. (2019). Thalamus Optimized Multi Atlas Segmentation (THOMAS): fast, fully automated segmentation of thalamic nuclei from structural MRI. Neuroimage, 194, 272–282.

Trzesniak, C., Linares, I. M., Coimbra, É. R., Júnior, A. V., Velasco, T. R., Santos, A. C., & Crippa, J. A. (2016). Adhesio interthalamica and cavum septum pellucidum in mesial temporal lobe epilepsy. Brain Imaging and Behavior, 10(3), 849–856.

Van der Linden, M., Adam, S., Agniel, A., Baisset-Mouly, C., Bardet, F., & Coyette, F. (2004). L’évaluation des troubles de la mémoire épisodique: fondements théoriques et méthodologiques. M. Van Der Linden, L’évaluation des troubles de la mémoire, Marseille, Solal, 11–20.

Van der Werf, Y. D., Scheltens, P., Lindeboom, J., Witter, M. P., Uylings, H. B., & Jolles, J. (2003). Deficits of memory, executive functioning and attention following infarction in the thalamus; a study of 22 cases with localised lesions. Neuropsychologia, 41(10), 1330–1344.

Van der Werf, Y. D., Witter, M. P., Uylings, H. B., & Jolles, J. (2000). Neuropsychology of infarctions in the thalamus: a review. Neuropsychologia, 38(5), 613–627.

Vidal, J. P., Danet, L., Péran, P., Pariente, J., Bach Cuadra, M., Zahr, N. M., … & Saranathan, M. (2024). Robust thalamic nuclei segmentation from T1-weighted MRI using polynomial intensity transformation. Brain Structure and Function, 229(5), 1087–1101.

Vidal, J. P., Rachita, K., Servais, A., Péran, P., Pariente, J., Bonneville, F., … & Barbeau, E. J. (2024b). Exploring the impact of the interthalamic adhesion on human cognition: insights from healthy subjects and thalamic stroke patients. Journal of Neurology, 271(9), 5985–5996.

Winkler, A. M., Ridgway, G. R., Webster, M. A., Smith, S. M., & Nichols, T. E. (2014). Permutation inference for the general linear model. Neuroimage, 92, 381–397.

Wong, A. K., Wolfson, D. I., Borghei, A., & Sani, S. (2021). Prevalence of the interthalamic adhesion in the human brain: a review of literature. Brain Structure and Function, 226, 2481– 2487.

Yarkoni, T., Poldrack, R. A., Nichols, T. E., Van Essen, D. C., & Wager, T. D. (2011). Large-scale automated synthesis of human functional neuroimaging data. Nature methods, 8(8), 665–670.

Yoneoka, Y., Takeda, N., Inoue, A., Ibuchi, Y., Kumagai, T., Sugai, T., … & Ueda, K. (2004). Acute Korsakoff syndrome following mammillothalamic tract infarction. American journal of neuroradiology, 25(6), 964–968.

